# Translational control by *Trypanosoma brucei* DRBD18 contributes to the maintenance of the procyclic state

**DOI:** 10.1101/2023.02.08.527724

**Authors:** Martin Ciganda, José Sotelo-Silveira, Joseph T. Smith, Shichen Shen, Jun Qu, Pablo Smircich, Laurie K. Read

## Abstract

*Trypanosoma brucei* occupies distinct niches throughout its life cycle, within both the mammalian and tsetse fly hosts. The immunological and biochemical complexity and variability of each of these environments require a reshaping of the protein landscape of the parasite both to evade surveillance and face changing metabolic demands. Whereas most well-studied organisms rely on transcriptional control as the main regulator of gene expression, post-transcriptional control mechanisms are particularly important in *T. brucei*, and these are often mediated by RNA-binding proteins. DRBD18 is a *T. brucei* RNA-binding protein that interacts with ribosomal proteins and translation factors. Here, we tested a role for DRBD18 in translational control. We show that DRBD18 depletion by RNA interference leads to altered polysomal profiles with a specific depletion of heavy polysomes. Ribosome profiling analysis reveals that 101 transcripts change in translational efficiency (TE) upon DRBD18 depletion: 41 exhibit decreased TE and 60 exhibit increased TE. A further 66 transcripts are buffered, *i.e*. changes in transcript abundance are compensated by changes in TE such that the total translational output is expected not to change. Proteomic analysis validates these data. In DRBD18-depleted cells, a cohort of transcripts that codes for procyclic form-specific proteins is translationally repressed while, conversely, transcripts that code for bloodstream form- and metacyclic form-specific proteins are translationally enhanced. These data suggest that DRBD18 contributes to the maintenance of the procyclic state through both positive and negative translational control of specific mRNAs.

## INTRODUCTION

Infectious organisms regulate crucial processes such as proliferation and generation of infective stages through coordinated changes in their protein repertoire in response to changes in nutrients, immune response of the host, and other environmental factors such as temperature (Verma-Gaur and Traven 2016; El Mouali and Balsalobre 2019; Johansson and Freitag 2019). *Trypanosoma brucei* is a parasite with a complex life cycle that involves a *Glossina* insect vector and a mammalian host, in which regulated differentiation proceeds through distinctly adapted life stages (Matthews 2005). Within the mammalian host, proliferation of slender bloodstream forms (BF) is accompanied by differentiation via quorum sensing mechanisms into cell cycle arrested stumpy forms that are preadapted to progress into insect forms once taken up in a blood meal (Quintana et al. 2021). Within the tsetse fly midgut, parasites differentiate into the procyclic form (PF) (Gunasekera et al. 2012). As nutrient availability and environmental conditions within the midgut change, PF parasites begin their migration towards the salivary glands, where infective metacyclic form are generated (Christiano et al. 2017).

In trypanosomes, control of gene expression is largely posttranscriptional (Clayton 2019), taking place through the action of *trans* acting factors on *cis* elements in the UTRs of the mRNAs (or occasionally, the coding regions) (Kramer and Carrington 2011). RNA-binding proteins (RBPs) are critical *trans* acting factors that control steady state levels of mRNAs by regulating mRNA processing, nuclear export, degradation, and translatability (Kolev et al. 2014; Clayton 2019). For example, the PuREBP1/2 complex, which contains two RBPs, controls the purine-dependent decay of the NT8 nucleobase transporter mRNA in PF *T. brucei* by binding a stem loop element its 3’ UTR (Rico-Jimenez et al. 2021). RNA-binding protein, Puf9, stabilizes target transcripts involved in kinetoplast replication during S-phase (Archer et al. 2009). RBPs have also emerged as fundamental players in the regulation of the *T. brucei* life cycle (Kolev et al. 2014; Clayton 2019). RBP10 is a BF-specific protein that binds a U(A)U_6_ motif in the 3’ UTR of many target RNAs and is required for growth of *T. brucei* as BFs (Wurst et al. 2012; Mugo and Clayton 2017). Cells depleted of RBP10 can only grow as PFs, and if PFs are stimulated to overexpress RBP10, they can only grow as BFs. RBP7 is one of over 30 molecules identified in response to quorum sensing that triggers the transition from slender BFs to quiescent stumpy BF (McDonald et al. 2018). Finally, overexpression of RBP6 in PF triggers their differentiation to MFs. Despite their critical importance in all facets of growth and development, most *T. brucei* RBPs remain uncharacterized.

DRBD18 is a double RRM-containing protein that is both essential and abundant, and whose depletion leads to a large remodeling of the transcriptional landscape in BF and PF *T. brucei* (Lott et al. 2015; Bishola Tshitenge and Clayton 2022). In addition, we previously showed that DRBD18 has a role in the nuclear export of a subset of mRNAs (Mishra et al. 2021). Subsequent work by others has demonstrated that depletion of DRBD18 leads to defects in pre-mRNA processing (Bishola Tshitenge and Clayton 2022). Thus, DRBD18 is a multifunctional RBP. Interestingly, DRBD18 was found associated with polysomes in BF cells (Klein et al. 2015), and DRBD18 pull downs recovered numerous ribosomal proteins and translation factors in BF and PF (Lott et al. 2015; Bishola Tshitenge and Clayton 2022), suggesting a direct role for DRBD18 in translational control. Multiomic approaches involving ribosome profiling (Ribo-seq) have been successfully used to address several aspects of translational control in trypanosomes (Jensen et al. 2014; Vasquez et al. 2014; Smircich et al. 2015; Chavez et al. 2021). Here, we use these approaches to explore the function of DRBD18 in translational regulation in PF *T. brucei*.

## RESULTS

### DRBD18 associates with ribosomes and translation initiation factor eIF3a

Previous studies from our laboratory and others demonstrated the presence of ribosomal proteins and translation factors among the proteins enriched by association with DRBD18 in *T. brucei* (Lott et al. 2015; Bishola Tshitenge and Clayton 2022) (**Table S1**), suggesting a role for DRBD18 in translation. To determine if DRBD18 is associated with translating ribosomes in PF, we analyzed sucrose gradient fractions from sedimented lysates from cycloheximide-treated cells (**Fig. 1A**). As shown by continuous measurement of absorbance at 260 nm (blue trace), fractions were collected containing ribosome-free components (RF), ribosomal subunits, monosomes (M), and polysomes. Western blot analysis demonstrated that DRBD18 can be detected in fractions from the polysome-enriched, dense portions of the gradients. To confirm that the presence of DRBD18 in these fractions correlates with the presence of polysomes, we treated control lysates with EDTA, disassembling ribosomes. As shown in the bottom panel, monosomes and polysomes are dissociated upon EDTA treatment, and concomitantly, DRBD18 is lost from the high-density fractions. Thus, a fraction of DRBD18 associates with monosomes and polysomes.

**Figure 1.**
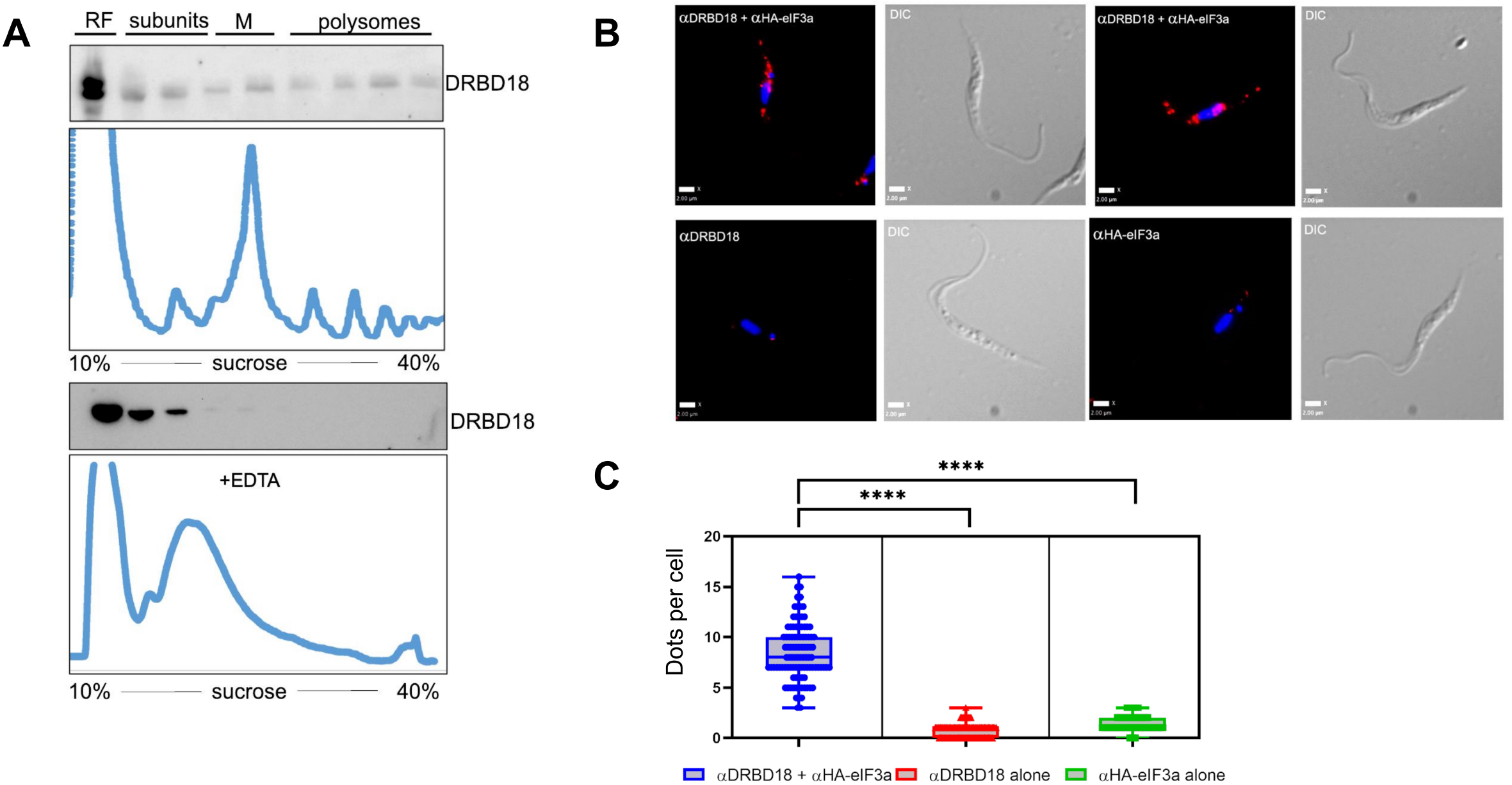
DRBD18 associates with polysomes and translation initiation factor eIF3a. (*A*) Anti-DRBD18 western blot detection of DRBD18 in high-density, polysomal fractions from cycloheximide-treated lysates separated on 10-40% sucrose gradients (top). EDTA-mediated dissociation of ribosomes leads to loss of DRBD18 association with high-density fractions (bottom). (*B*) Proximity ligation assay in cells with C-terminally HA-tagged eIF3a using antibodies against HA and DRBD18. Two examples are shown in the top row with corresponding DIC images. Red dots indicate signal from amplification of individual interactions. Negative controls in the bottom row are individual antibodies alone; anti-DRBD18 (left), anti-HA (right), with corresponding DIC images. Bars = 2 microns (*C*) Quantification of dots per cell (100 cells per condition); both antibodies, blue; anti-DRBD18 alone, red; anti-HA alone, green. Students t-test, **** < 0.0001.

We previously identified nine subunits of the translation initiation factor, eIF3, in DRBD18 pull downs (**Table S1**). In addition to its well-characterized role in translation initiation, eIF3 can also be a negative regulator of translation, with some functions being subunit-selective (Valasek et al. 2017; Wolf et al. 2020). In *T. brucei*, eIF3 is composed of 11 subunits (subunits a through l, with the exception of subunit j), and they can be tagged, overexpressed and co-purified in a complex (Li et al. 2017). To define the association of DRBD18 and the eIF3 complex, we selected eIF3a for further study as this subunit was also detected in association with DRBD18 in BF (Bishola Tshitenge and Clayton 2022). To achieve high resolution and *in situ* specificity, we utilized a proximity ligation assay (PLA) (Fredriksson et al. 2002) to visualize the interaction between DRBD18 and eIF3a-HA in fixed and permeabilized cells. When candidate interactors are within 40 nm, the ligation and amplification reactions amplify this single event into a detectable fluorescence signal. As seen in **Fig. 1B**, when PF *T. brucei* are incubated with both □-DRBD18 and □-HA antibodies, fluorescence signals from the interactions (artificially colored in red) are observed (8 dots per cell, range 3 to 17; **Fig. 1C**), indicating that DRBD18 and eIF3a-HA are in close proximity, either interacting directly or as part of a complex. DRBD18 has been described as being cytoplasmic and nuclear, and it is likely that it shuttles between both compartments (Lott et al. 2015; Dean et al. 2017; Bishola Tshitenge and Clayton 2022). In turn, *T. brucei* eIF3 is cytoplasmic, consistent with its role in translation (Li et al. 2017). Topologically, therefore, the cytoplasm provides the most opportunities for interaction between eIF3a and DRBD18, and that is indeed what we observed. As controls, when individual primary antibodies are utilized alone, the number of dots per cell falls significantly (range 0-4, one hundred cells per condition quantified in the right-most panel; **Fig. 1C**). These data confirm specificity of the DRBD18-eIF3a interaction, which may be in the context of the canonical eIF3 complex or within a distinct complex. Interaction of DRBD18 with both translating ribosomes and translation initiation factor, eIF3a, supports a role for DRBD18 in translation.

### Depletion of DRBD18 leads to a translational defect

Having shown a physical association between DRBD18 and components of the translational machinery, we next investigated the global translational status of PF cells under conditions of RNAi-mediated DRBD18 depletion. Lysates from uninduced and induced cells were layered onto sucrose gradients and separated by ultracentrifugation so that ribosomal subunits, monosomes, and polysomes could be discriminated and visualized. Integrating the area under each peak gives a readout of total mass in the selected category, and the monosome/polysome area ratio provides an indication of translational status (Rowe et al. 2014; Chassé et al. 2017): the lower this ratio, the more ribosome mass will be engaged in higher density polysomal fractions actively translating mRNAs. Conversely, the higher the ratio, the less ribosomal mass will be engaged in higher translational output. After induction of DRBD18 RNAi, the monosome/polysome ratio is increased (**Fig. 2A**), reflecting a decrease in translational activity mediated by DRBD18 depletion. Four replicates of the experiment were quantified, and the monosome/polysome ratios are shown in **Fig. 2B**. To rule out the possibility that this phenotype is a consequence of indirect stress-related processes, we analyzed polysomal profiles of PF cells depleted of the essential RBP, RBP16 (Pelletier and Read 2003). We performed the assays at 24 hours post-induction, a time point at which DRBD18 and RBP16 cell lines both exhibit growth phenotypes (**Fig. S1**). The decrease in polysomes in the DRBD18-depleted cells, but not in the RBP16-depleted cells (**Fig. S1**), demonstrates that DRBD18 knockdown leads to a specific decrease in translating polysomes.

**Figure 2.**
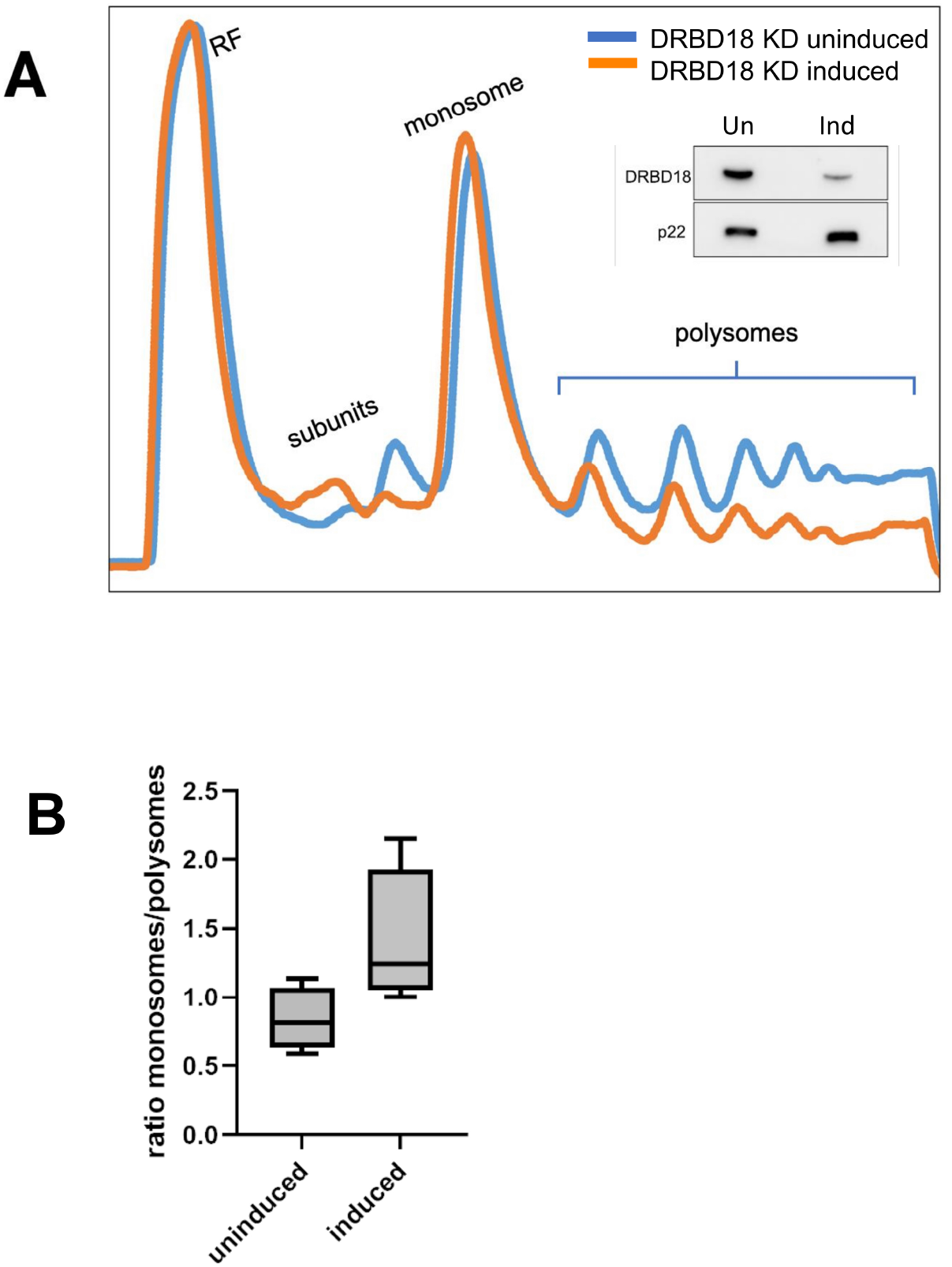
Depletion of DRBD18 in procyclic cells leads to translational defects. (*A*) Polysomal profiles of lysates from PF DRBD18 RNAi cells either uninduced (blue) or doxycycline-induced (red). RF; ribosome free fractions. Inset: Western blot of induced and uninduced whole cell samples showing efficient knockdown of DRBD18; p22 is used as a loading control. (*B)* Quantification of the monosome to polysome ratio from four independent experiments as in (*A*).

### Depletion of DRBD18 leads to changes in the translational efficiencies of a subset of transcripts

To gain a better understanding of how translational regulation through DRBD18 occurs in PF *T. brucei*, we performed ribosome profiling comparing DRBD18-depleted cells (induced with doxycycline) to DRBD18 replete cells (uninduced). The length distribution of reads was consistent with *bona fide* ribosomal footprints (**Fig. S2**) (Ingolia et al. 2009; Jensen et al. 2014; Vasquez et al. 2014; Smircich et al. 2015). Analysis using the anota2seq package (Oertlin et al. 2022) revealed that 102 transcripts were significantly altered in their translational efficiencies, resulting in either an increase (41 transcripts, TE UP) or a decrease (61 transcripts, TE DOWN) in translation efficiency (**Figure 3**, light and dark red, respectively; **Tables S2 and S3**). Note that because the transcript for DRBD18 was targeted via RNA interference using the 3’ UTR, many reads were counted for the gene in the induced samples, artificially marking DRBD18 as being reduced in translation. After a closer analysis of read distribution in all the samples, DRBD18 was consequently removed from the list of affected genes, leaving 60 transcripts in the TE DOWN set and a total of 101 transcripts with altered translational efficiency that is expected to lead to increased protein levels. Additionally, we identified another set of transcripts for which the rates of association with ribosomes change when DRBD18 is depleted, but because the transcript abundance changes significantly in the opposite direction their translational output is buffered in a way that no overall changes in protein levels are expected (**Table S3**). These are labeled “buffered”, and colored light and dark blue in **Fig. 3**. A final set of transcripts is changed at the level of mRNA abundance, which is accompanied by the expected change in translation rates (thus, no change in TE is observed) (**Fig. 3**, green; **Table S3**). Given the level of translational repression observed upon DRBD18 knockdown in **Fig. 2**, it was somewhat surprising that only 60 mRNAs were identified as TE DOWN. However, some of the transcripts that are translated less efficiently, and therefore lose association with the ribosome, are present in high abundance (*e.g*., EP1 procyclin, **Fig. S3**), and their loss from higher order polysomes may account for the observed phenotype. Overall, these data demonstrate that DRBD18 impacts the translational efficiency of numerous mRNAs both positively and negatively.

**Figure 3.**
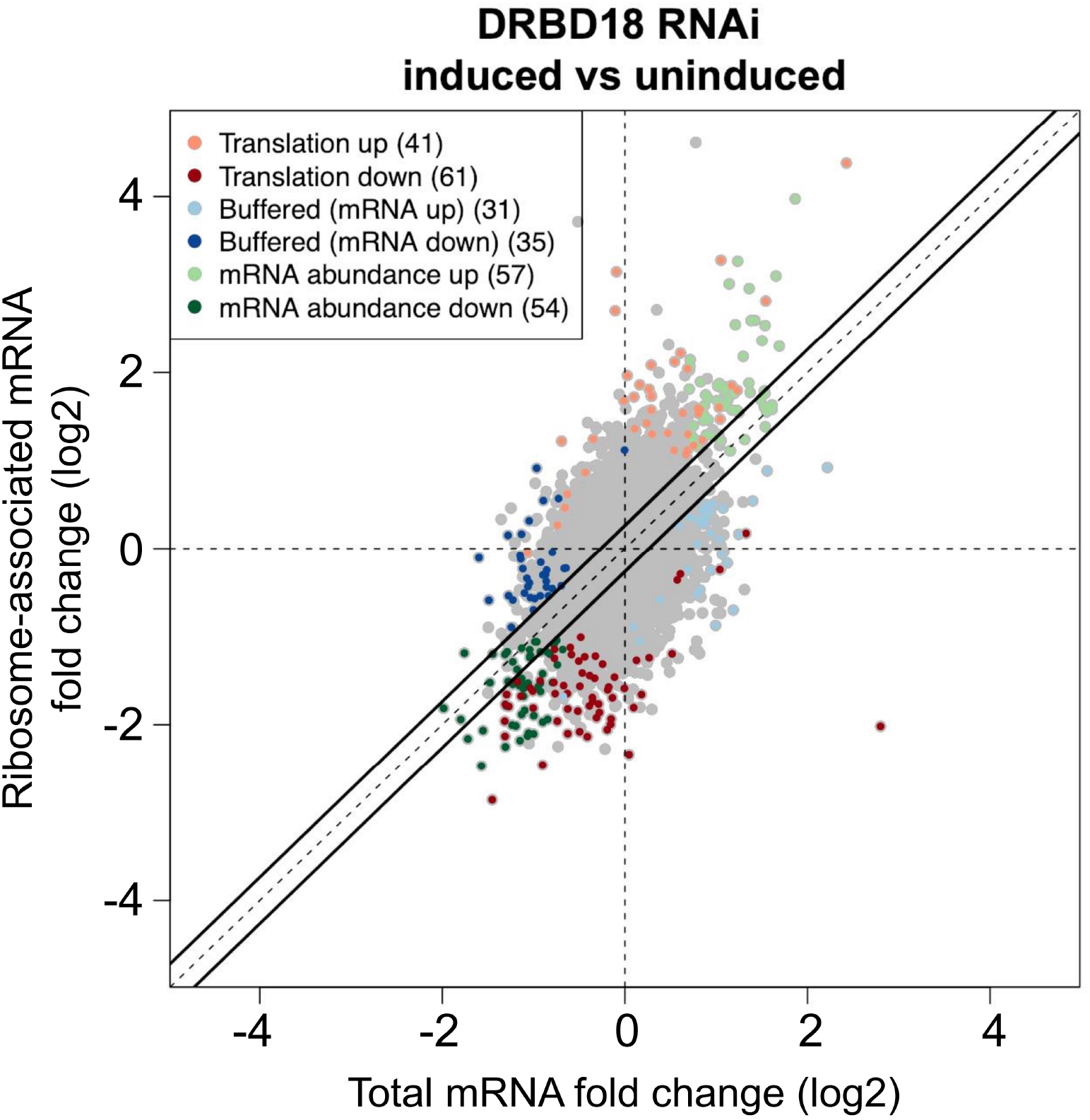
Ribosome profiling of cells depleted of DRBD18 reveals changes in translational efficiency. Anota2seq plot showing changes ribosome-associated transcripts against changes in abundance of total transcripts. mRNAs with altered translational efficiency (Translation), either enhanced or repressed, are in light and dark red, respectively. mRNAs that change at the level of abundance, but are then buffered at the translational level so that their protein output is not expected to change (Buffered), either in transcripts with lower or higher abundance, are in light and dark blue, respectively. mRNAs that are regulated mainly by their abundance (Abundance), either upregulated or downregulated, are in light and dark green, respectively.

To begin to characterize the set of mRNAs that are translationally regulated by DRBD18, we analyzed GO terms enriched in these sets. In the TE UP genes, enriched GO categories include those involved in C-terminal protein methylation, signal transduction, mRNA processing, mRNA cleavage, and posttranslational modifications; however, none of these reached significance after Bonferroni correction. For the genes enriched in the TE DOWN set, we detected terms related primarily to adenylate cyclase activity (**Fig. 4A**). Indeed, seven adenylate cyclases exhibited decreased TE upon DRBD18 depletion (**Tables S2 and S3**), suggesting that cAMP metabolism may be altered in DRBD18 knockdown cells.

**Figure 4.**
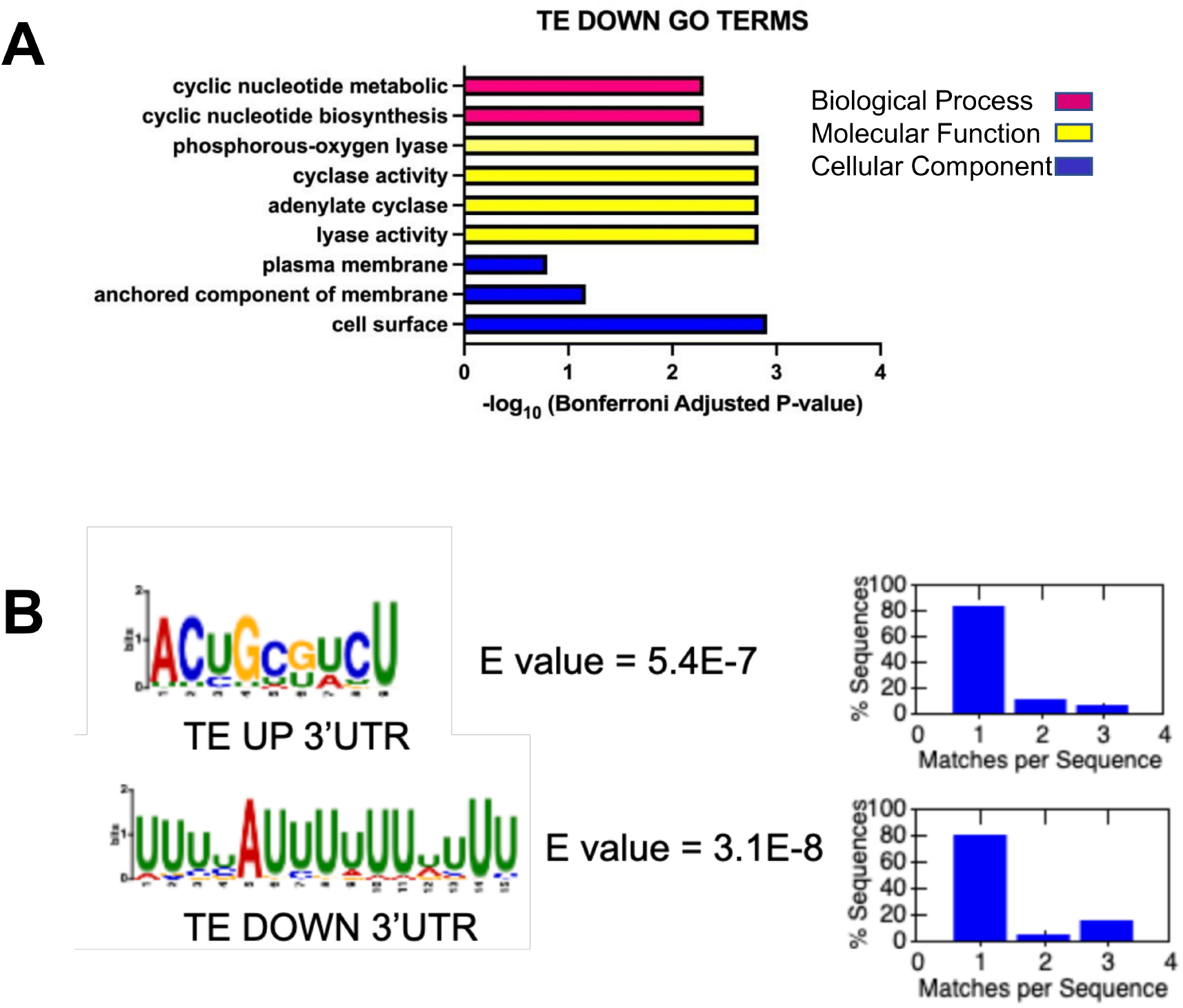
GO term and motif analysis of translationally regulated mRNAs. (*A*) Enriched GO term categories for the TE DOWN mRNAs are indicated on the left with the –log_10_ (Bonferroni adjusted P-value) shown on the bottom. Red bars correspond to enriched biological process categories, yellow bars to enriched molecular functions categories, and blue bars correspond to categories enriched cellular component categories. TE UP genes were not significant after Bonferroni adjustment. (*B*) Enriched motifs in the 3’ UTRs of TE UP and TE DOWN sets. XSTREME analysis was performed on annotated 3’ UTRs longer than 6 nucleotides. Sequence logo plot is illustrated for the most significant matches in the TE UP and TE DOWN datasets, with significance and number of motifs per sequence indicated to the right.

As translational regulatory elements are often found in 3’ UTRs, we next determined whether the TE UP and TE DOWN transcripts contain conserved motifs in their 3’ UTRs using XSTREME (Grant 2021). Of the 41 genes with increased TE and 60 genes with decreased TE, 26 (63%) and 38 (63%), respectively, had annotated 3’UTRs that allowed comprehensive motif analysis. These UTRs had varying lengths (range 75 nt to 11,187 nt for those in the TE UP set, averaging 2038 nt; range of 38 to 4007 nt for the TE DOWN set, averaging 693 nt). A search for motifs using standard parameters (length 6 to 15 expected anywhere in the sequence) yielded six and eight motifs, respectively. **Fig. 4B** shows the most significant motifs by E value (by at least an order of magnitude): a 9 nt motif (ACUGCGUCU) for the TE UP dataset and a U-rich sequence with one conserved A residue for the TE DOWN dataset. Eighteen sites were discovered for this 9 nt motif in the TE UP dataset (present in 69.2% of input UTRs) and twenty of the U-rich sites were discovered in the TE DOWN dataset (52% of input UTRs). Both motifs were typically present in one copy per transcript (**Fig. 4B**). The distinct motifs found in the TE UP and TE DOWN transcript sets suggests these motifs could be involved in translational regulation.

To gain insight into possible mechanisms of action of DRBD18 in translation regulation, we asked whether DRBD18 reportedly binds the translationally regulated mRNAs. To this end, we took advantage of an available DRBD18 RIP-seq dataset (Bishola Tshitenge and Clayton 2022). It should be kept in mind that we interpret these results cautiously because this dataset was obtained in BF, whereas our ribosome profiling dataset was obtained in PF. Nonetheless, the transcripts we analyze are also expressed in PF, so the assumption that they are bound by DRBD18 in PF is not unreasonable. We analyzed our TE UP and TE DOWN transcripts in the context of the 235 transcripts reported to bind with high confidence to DRBD18 in BF (2-fold eluate/flow through and padj<0.05) (**Fig. 5A**). By plotting the eluate/flow through ratios, we identified six bound transcripts that are also translationally regulated by the depletion of DRBD18 in PF, three that are TE UP and three TE DOWN (**Fig. 5B**). For example, the adenylate cyclase, ACP6, a negative regulator of social motility (SoMo) (Lopez et al. 2015; Shaw et al. 2022), is translationally repressed in DRBD18 knockdowns. Conversely, PuReBP2, a component of the complex that represses NT8 nucleobase transporter mRNA abundance in a purine-responsive manner (Rico-Jimenez et al. 2021), is translationally enhanced upon DRBD18 knockdown. Together, these six transcripts are candidates for direct translational regulation by DRBD18.

**Figure 5.**
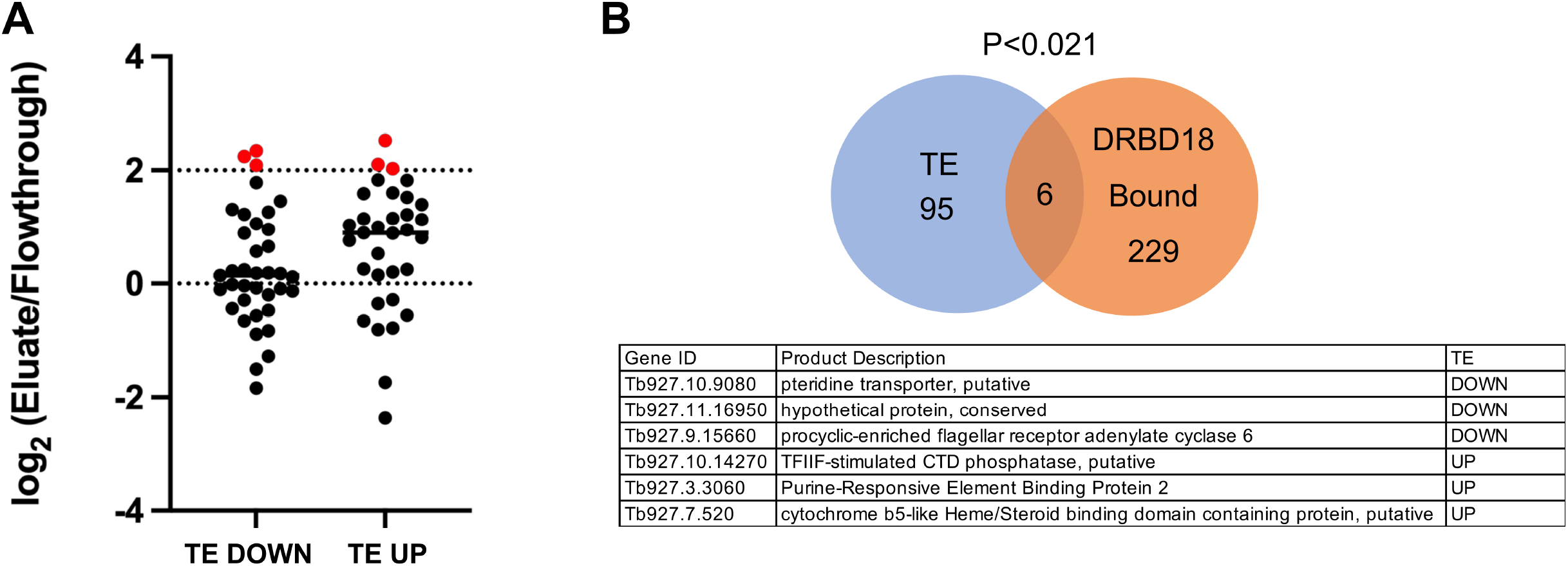
A subset of transcripts regulated at the level of translational efficiency by DRBD18 also associate with DRBD18. *(A)* Binding values (expressed as -log of ratios between normalized reads associated with eluate and those in the unbound fraction) from a published dataset (Bishola Tshitenge and Clayton 2022) were plotted for all the transcripts in the TE categories. A stringent cutoff value of 2 was selected to indicate significant binding. Transcripts meeting this threshold are indicated in red. *(B)* Venn diagram representation of intersection between the datasets and hypergeometric test results indicating significance. The list of intersecting genes is given below.

### DRBD18 regulates translational efficiency of life cycle stage-specific transcripts

To further assess the impact of DRBD18-mediated translational regulation, we asked whether there were any life cycle stage-specific patterns to the translationally affected transcripts. To this end, we took advantage of published datasets that describe protein expression changes during the parasite life cycle. We used a tool to visualize proteome-wide ordered protein abundances in BF *vs*. PF cells (Tinti and Ferguson 2022) to highlight the TE UP and TE DOWN sets (**Fig. 6A**, magenta and yellow, respectively). In this published proteome, we identified 26 genes from our TE UP and 42 genes from our TE DOWN datasets. Of these, we observed a correlation between transcripts in the TE DOWN set and those proteins that are more highly expressed in PF (**Fig. 6A, vertical box**). Similarly, we identified a correlation between transcripts in the TE UP set and those proteins that are more highly expressed in BF (**Fig. 6A, horizontal box**). To quantify these associations, we obtained intersection lists between the TE UP and TE DOWN transcripts and the PF- and BF-specific proteins (defined arbitrarily by manually selecting values below 3.7 on a given axis), and we tested these lists by hypergeometric tests to determine the significance of the overlap. For TE UP and PF-specific proteins, and TE DOWN and BF-specific proteins, the overlaps were not statistically significant. However, for TE DOWN and PF-specific proteins, a significant overlap was observed (**Fig. 6B**). This set included numerous adenylate cyclase gene products, and two transcripts that are reportedly bound by DRBD18 (**Fig. 6B**). While the TE UP and BF-specific proteins overlap did not reach statistical significance due to the large number of BF proteins in the published proteome, 41% (12/29) of the TE UP genes that were also present in the proteome are BF-specific (**Fig. 6C**). Interestingly, in the set of TE UP transcripts whose protein products are BF-specific, we identified RBP10, a positive regulator of the BF life cycle stage (Mugo and Clayton 2017). To confirm this finding, we performed western blot analysis of RBP10 in DRBD18 replete and depleted conditions. As expected, almost no RBP10 is detected in uninduced PF DRBD18 RNAi cells (**Fig. 6D**). However, when DRBD18 is depleted, a dramatic increase we observe a RBP10 signal, identifying DRBD18 as a negative regulator of RBP10 protein production in PF. Overall, these data demonstrate that DRBD18 plays a role in maintaining the PF state by enhancing translation of a subset of PF-specific proteins and repressing translation of a subset of BF-specific proteins.

**Figure 6.**
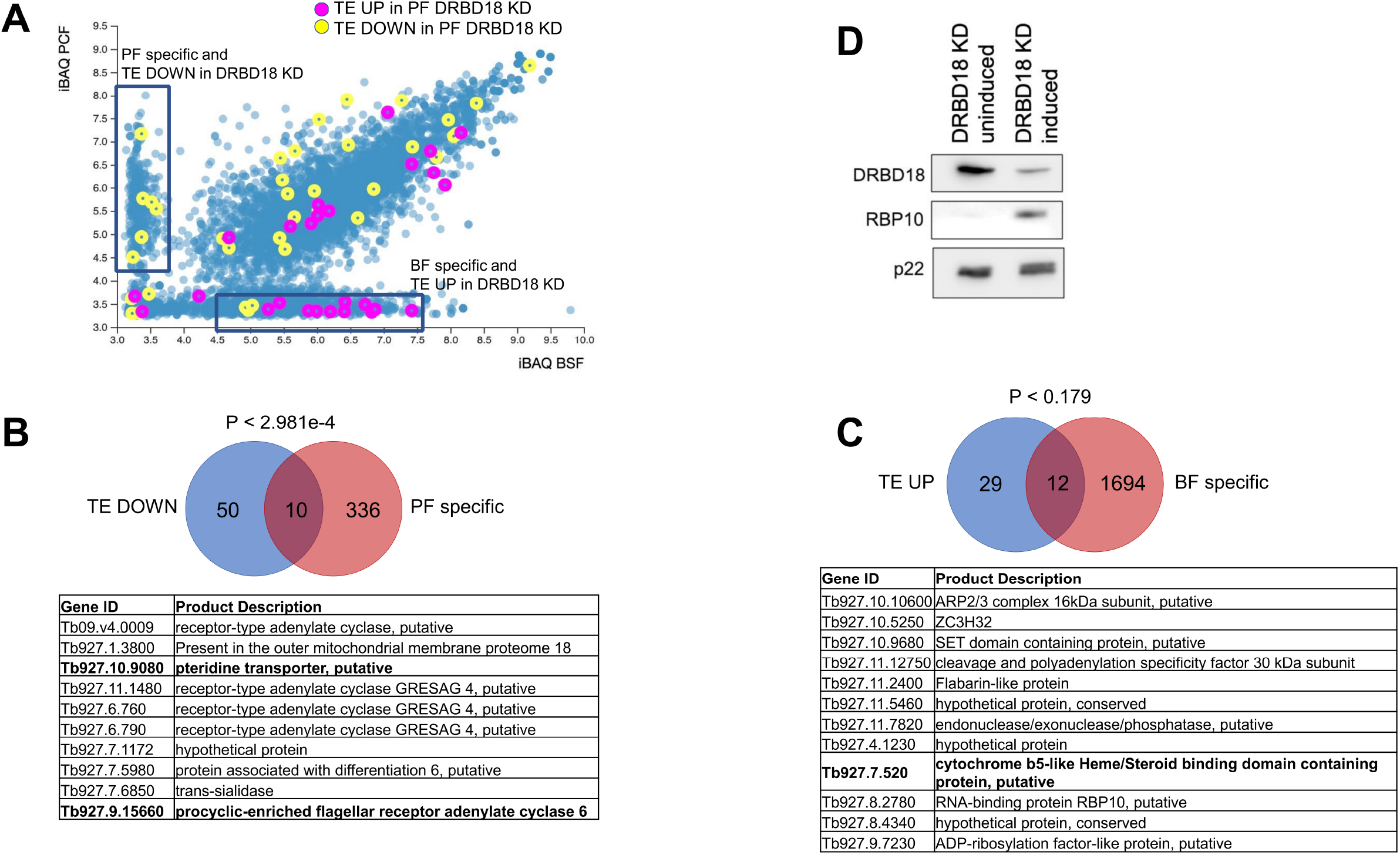
DRBD18 contributes to the maintenance of the procyclic translatome by repressing translation of bloodstream-specific and enhancing translation of procyclic-specific transcripts. *(A)* An available dataset of protein abundances (Tinti and Ferguson 2022) in procyclic *vs*. bloodstream life cycle stages is plotted with genes from the TE UP and TE DOWN categories highlighted in magenta and yellow, respectively. A cutoff value of 3.7 on each axis was arbitrarily selected to enrich protein lists in procyclic- and bloodstream-specific proteins. *(B)* and *(C)* Venn diagrams showing intersections of TE DOWN and TE UP categories with procyclic- and bloodstream-specific proteins, respectively. Hypergeometric tests were performed to assess significance of these intersections. *(B)* The list of TE DOWN genes that are procyclic-specific is shown. Those that are also bound by DRBD18 in BF (Bishola Tshitenge and Clayton 2022) (Fig. 5) are bolded. *(C)* The list of TE UP genes that are bloodstream-specific is presented as in *(B). (D)* Western blot of RPB10 (Tb927.8.2780), confirming its upregulation upon DRBD18 knockdown in procyclic forms. P22 is a loading control.

Given the life cycle-specific nature of many transcripts that are translationally regulated by DRBD18, we next asked if DRBD18 is also involved in translational regulation of MF-specific genes. In a previous study, mass spectrometry analysis identified sets of proteins whose abundance increases or decreases in MFs generated in culture by RPB6 overexpression (Christiano et al. 2017). We first compared our translationally DRBD18-regulated datasets to the MF/PF protein expression ratios from this study (**Fig. 7A**). We identified a significant difference between the TE DOWN and TE UP transcripts with regard to their expression in MF vs. PF (**Fig. 7**). Transcripts that are translationally repressed in the absence of DRBD18 tend to have low relative MF/PF expression. Conversely, transcripts that are translationally enhanced in the absence of DRBD18 tend to have higher MF expression relative to PF expression. We next compared the gene lists from our TE UP dataset and those transcripts that are increased in MF (MF up) and identified seven overlapping genes; this overlap was significant as indicated by a hypergeometric test (**Fig. 7B**). Similarly, comparison of the overlap between our TE DOWN dataset and those genes that are decreased in MF identified nine overlapping genes, again with a significant overlap (**Fig. 7C**). Thus, DRBD18 further contributes to maintenance of the PF state by repressing translation of a set of MF-specific transcripts and enhancing translation of transcripts that are more highly expressed in PF.

**Figure 7.**
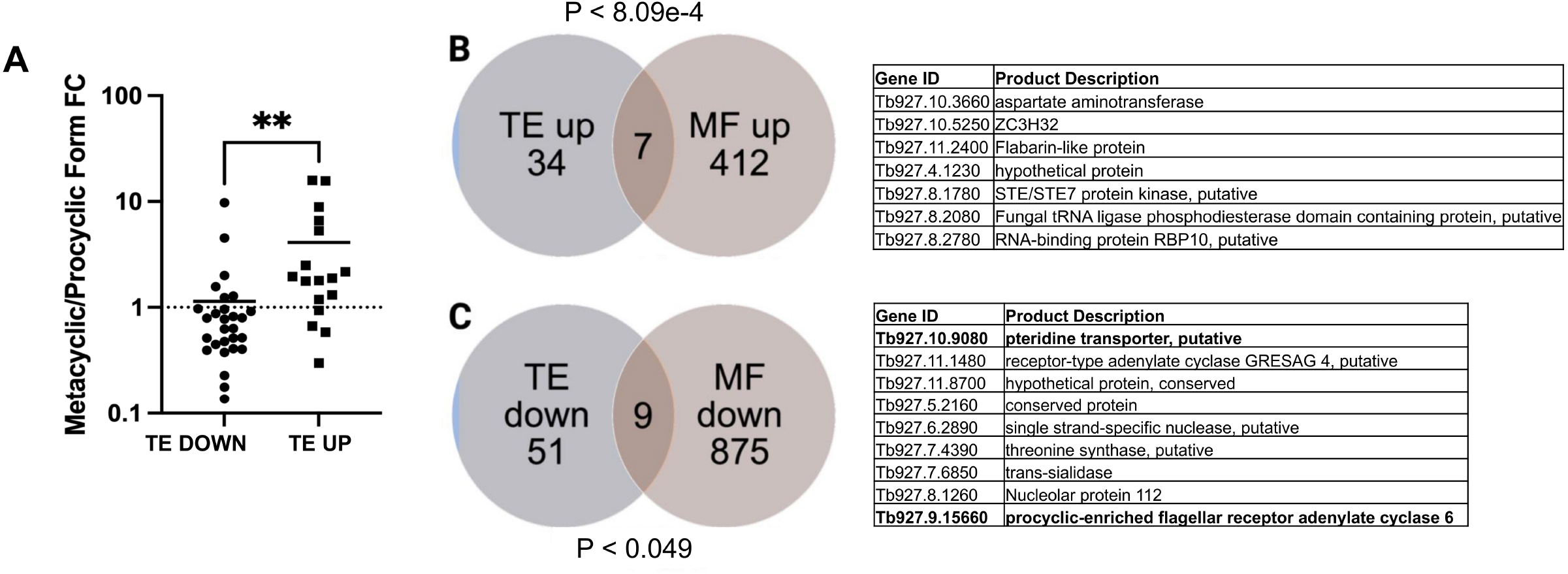
DRBD18 contributes to the maintenance of the procyclic translatome by repressing translation of metacyclic-specific and enhancing translation of procyclic-specific transcripts. *(A)* An available dataset of protein abundances in procyclic versus metacyclic life cycle stages (Christiano et al. 2017) was analyzed for correlation with our TE dataset. Transcripts were segregated by TE DOWN or TE UP, and the protein fold change (FC) in metacyclics *vs*. procyclics was plotted for each gene that was present in the proteome.,**; p = .0063 by Student’s t-test. *(B)* Venn diagram showing intersection of TE UP and metacyclic (MF) up genes; identities of the seven overlapping genes are shown to the right. *(C)* Venn diagram showing the intersection of TE DOWN and MF down genes; identities of the nine overlapping genes are shown to the right. In *(B)* and *(C)*, hypergeometric tests were performed to assess significance of these intersections.

### Global proteome of PF cells depleted in DRBD18 reveals that protein level changes accompany changes in TE

Finally, we asked whether we could identify proteomic changes in DRBD18-depleted cells consistent with the translation effects identified by ribosome profiling. Using quantitative mass spectrometry, we detected 4,734 proteins with an average of 8.1 peptides per protein (**Table S4**). We identified 322 proteins that yielded an >50% fold change upon DRBD18 RNAi (protein ratio <0.67 or >1.5) while reaching a level of significance (P < 0.05). Of these, 152 are upregulated, and 170 are downregulated. We identify by mass spectrometry peptides corresponding to 23 out of the 41 TE UP transcripts. Of these 23, eight were upregulated at the protein level (**Fig. 8; Table S4**). As expected, RBP10 (Tb927.8.2780) was significantly upregulated. In addition, ZC3H32 (Tb927.10.5250), a BF-specific, essential cytosolic mRNA-binding protein that can repress translation when tethered to a reporter (Klein et al. 2017), is also upregulated. Glycogen synthase kinase 3 (Tb927.10.13780), an enzyme previously studied as a potential drug target (Ojo et al. 2008), was upregulated upon DRBD18 knockdown as well. For the 60 TE DOWN transcripts, we identified corresponding peptides for 28 out of these 60 by mass spectrometry, and eight of the corresponding gene products were identified in the downregulated proteome (**Fig. 8; Table S4**). These include GRESAG4 (Tb927.11.1480), an adenylate cyclase that is translationally enhanced PF (Durante et al. 2020). The absence of corresponding changes in the set of translationally regulated mRNAs and our global proteome data may have several explanations. For example, we noted that some proteins encoded by TE UP and TE DOWN transcripts did change abundance in the expected direction, but were slightly outside our significance criteria. For others, this discrepancy may be explained by the existence of post-translational controls, such as phosphorylation, methylation, or ubiquitination that modulate protein stability. Collectively, these data demonstrate that DRBD18 impacts the PF *T. brucei* proteome, and does so at least in part by affecting the translational efficiency of a subset of transcripts.

**Figure 8.**
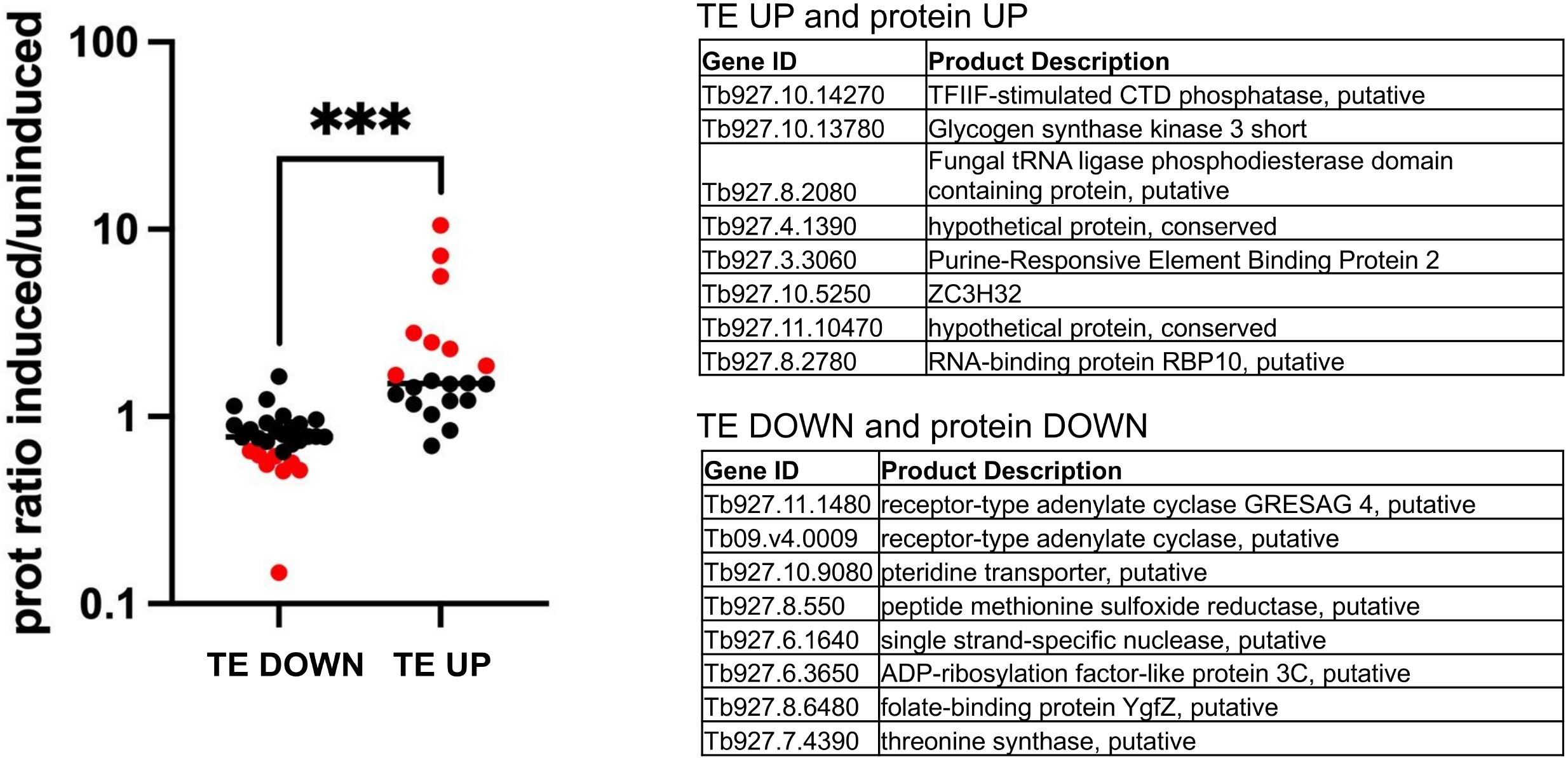
Changes in the proteome accompany TE changes detected by ribosome profiling in DRBD18-depleted cells. The protein ratio in cells depleted of DRBD18 (induced) *vs*. that in uninduced cells is plotted for our TE DOWN and TE UP datasets. The significance between the TE DOWN and TE UP datasets was calculated by Student’s t-test; *** indicates p = 0.0007. The TE genes whose products are also significantly altered in the proteome are indicated in red and listed on the right.

## DISCUSSION

Translational control constitutes a major mechanism of gene regulation in *T. brucei*, including during life cycle progression (Jensen et al. 2014; Vasquez et al. 2014). However, the factors that impact translatability of specific transcripts are poorly understood. In this manuscript, we report an expanded gene regulatory role for the *T. brucei* RNA binding protein, DRBD18, using ribosome profiling to quantify its impact on the TE of PF mRNAs. We found that DRBD18 regulates the translational outputs of over 100 mRNAs in PF cells, with both increases and decreases in TE observed upon DRBD18 knockdown. Depletion of DRBD18 shifts the translational landscape away from the prototypical PF state, decreasing TE of PF-specific mRNAs and increasing the TE of both BF- and MF-specific mRNAs, highlighting a role for DRBD18-mediated translational control in life cycle stage maintenance.

The impacts of DRBD18 on translation could be direct or indirect, with specific mRNAs subject to distinct mechanisms. In support of a direct role for DRBD18 in modulating translation of some mRNAs, we show here that a fraction of DRBD18 associates with translating ribosomes. It is not yet known if this association promotes or inhibits translation of DRBD18-bound transcripts, or whether there are transcript-specific effects. Additionally, we show that DRBD18 interacts with the translation initiation factor, eIF3a. Our previous pulldown/mass spectrometry study identified the majority of *T. brucei* eIF3 subunits with DRBD18 in PF, suggesting DRBD18 interacts with the intact eIF3 complex (Lott et al. 2015). However, a similar study in BF *T. brucei* revealed only the eIF3a subunit (Bishola Tshitenge and Clayton 2022). These findings may reflect life cycle-specific differences in DRBD18 interactions, or they may be due to technical differences. eIF3a pulldowns in PF *T. brucei* did not identify DRBD18 (Li et al. 2017), suggesting that only a small subset of total eIF3a binds DRBD18. Beyond its well-known function in assembly of the 43S preinitiation complex, eIF3 has recently been reported to function in all stages of translation, including non-canonical modes of translation initiation, repression, and termination, as well as nonsense mediated decay and protein quality control, with some functions being subunit selective (Valasek et al. 2017; Wolf et al. 2020). The eIF3a subunit reportedly exerts both positive and negative transcript specific effects on translation in mammalian cells. Thus, it will be exciting in the future to determine the precise interactions between DRBD18 and eIF3 and how these modulate the translation of specific transcripts.

There is also substantial potential for DRBD18 to have an indirect role in mediating TE of distinct mRNAs. While DRBD18 impacts nuclear export of a subset of mRNAs (Mishra et al. 2021), these effects cannot be contributing to the TE changes seen here as we measured polysome-associated mRNA compared to cytoplasmic mRNA (see Materials and Methods). More importantly, DRBD18 substantially impacts 3’UTR length by modulating poly(A) addition sites in both BF and PF *T. brucei* (Bishola Tshitenge and Clayton 2022) (Tylec and Read, unpublished). Utilization of differential poly(A) addition sites on a given mRNA presumably leads to inclusion and exclusion of translational regulatory elements in mRNA 3’UTRs (Clayton 2019). Current experiments are focused on identifying the cohort of DRBD18-mediated changes in poly(A) addition sites in PF. These studies will then allow us to define the intersection between mRNAs with altered TE in this study and those with altered 3’ UTRs, and to define and test potential translational regulatory elements.

Comparison of our dataset of mRNAs that are translationally regulated by DRBD18 with published datasets of life cycle-specific protein expression (Christiano et al. 2017; Tinti and Ferguson 2022) uncovered a previously unknown and important role for DRBD18: contributing to the maintenance of the PF state. DRBD18 knockdown led to derepressed translation in PF of numerous transcripts that are normally upregulated in BF and MF. One of the most striking was RBP10, a BF specific repressor that strongly promotes the BF life cycle stage in *T. brucei* (Wurst et al. 2012; De Pablos et al. 2017; Mugo and Clayton 2017). Here, we show by ribosome profiling, mass spectrometry, and western blot that RBP10 translational output is increased after DRBD18 depletion in PF. In BF, RBP10 binds PF-specific mRNAs via a UA(U)_6_ motif, and targets them for degradation and translational inhibition, although the order of events is unclear (Mugo and Clayton 2017; Clayton 2019). Conversely, ectopic expression of RBP10 in PF, similar to the increased RBP10 expression we observe in PF after DRBD18 knockdown, led to cells that could only grow as BF and in which polysomal mRNA analysis suggested conversion to BF (Mugo and Clayton 2017). We show here that DRBD18 is critical for repressing RBP10 levels in PF, normally keeping both RBP10 mRNA abundance and TE low (**Tables S2 and S3**). Thus, we predict that one mechanism by which DRBD18 helps maintain the PF state is through its effect on RBP10 expression. Consistent with this prediction, the RBP10-binding motif, UA(U)_6_ is reminiscent of the top motif identified in our TE down dataset, suggesting that increased RBP10 may bind and contribute to translational repression of some PF-specific mRNAs upon knockdown of DRBD18 in PF. However, additional mechanisms must be at play, since many transcripts whose TE is altered in our dataset are not targets of RBP10, including 6 of the 10 that overlap the PF-specific set (Mugo and Clayton 2017). Another BF-specific protein whose TE is increased and which is upregulated in the proteome after DRBD18 knockdown is the RNA-binding protein, ZC3H32. ZC3H32 is a repressor that negatively impacts expression of reporter mRNAs to which it is tethered (Klein et al. 2017); thus, it may be repressing specific mRNA abundance and/or translation in PF DRBD18 knockdown cells. ZC3H32 is also upregulated 16-fold during metacyclogenesis (Christiano et al. 2017). Together, our data indicate that in PF *T. brucei*, DRBD18 normally acts upstream of both RBP10 and ZC3H32, repressing both their RNA abundance and translatability. This, in turn, keeps RBP10 and ZC3H32 from repressing specific PF transcripts, some of which may be represented in our TE DOWN dataset in DRBD18 knockdown cells.

The role of DRBD18 in repressing RBP10 expression in PF is somewhat unexpected, as DRBD18 plays the opposite role in BF. We show here that DRBD18 depletion leads to a 10-fold increase in RBP10 protein (**Table S4**), whereas in BF, DRBD18 depletion caused a 3-fold decrease in RBP10 protein (Bishola Tshitenge and Clayton 2022). One potential mechanism by which DRBD18 may effect such functions is its presence in distinct mRNPs. For example, two abundant, constitutively expressed *T. brucei* RBPs are DRBD2 and ZC3H41 (Clayton 2019). Of these, DRBD18 reportedly interacts with DRBD2 in BF and ZC3H41 in PF (Lott et al. 2015; Bishola Tshitenge and Clayton 2022), and distinct mRNPs likely have different roles in gene regulation. The basis for this binding specificity is unknown. However, arginine methylation of DRBD18 affects its protein-protein interactions in PF (Lott et al. 2015), and it is possible that the protein is differentially methylated in a life cycle specific manner. Together, these data implicate DRBD18 in promoting the PF state in PF, while promoting the BF state in BF.

With regard to stage-specific transcripts whose translation is normally promoted by DRBD18 in PF (those that are represented in the TE DOWN dataset), a major overrepresented class is PF specific adenylate cyclases, also called GRESAG4 isoforms, which represent five of the 10 transcripts in the TE DOWN/PF-specific dataset (**Fig. 6B**). GRESAG4 isoforms are an expanded family of <u>g</u>enes related to ESAG4 (<u>e</u>xpression <u>s</u>ite-<u>a</u>ssociated <u>g</u>ene 4) thought to compensate for the lack of canonical G protein-coupled receptors and heterotrimeric G proteins in *T. brucei* (Salmon 2018). Numerous members of this family are localized to the flagella, and the cAMP they produce functions in SoMo, chemotaxis, and migration of trypanosomes through their insect hosts. One GRESAG4 that we identified as being translationally enhanced by DRBD18 is ACP6 (Tb927.9.15660), a negative regulator of SoMo (Lopez et al. 2015; Shaw et al. 2022). We also identified Tb09.v4.0009, which is closely related to ACP6 and Tb927.11.1480 that shares with ACP6 localization to the flagellar tip (Lopez et al. 2015; Durante et al. 2020). The DRBD18-mediated positive regulation of translation of these GRESAG4 isoforms in PF suggests that DRBD18 impacts cell signaling through cAMP pathways with an effect on parasite motility. Interestingly, of the other two GRESAG4 isoforms identified in the set of transcripts that are TE DOWN upon DRBD18 knockdown, Tb927.6.760 and Tb927.6.790, the former is reportedly mitochondrially localized (Dean et al. 2017).

The current study has expanded our understanding of the critical regulatory RBP, DRBD18, in *T. brucei*. In addition to its previously described roles in regulating transcript abundance (Lott et al. 2015; Bishola Tshitenge and Clayton 2022), poly(A) site selection (Bishola Tshitenge and Clayton 2022), and mRNA nuclear export (Mishra et al. 2021), we show here that transcript-specific regulation of translational efficiency contributes to DRBD18’s remodeling of the PF proteome. Overall, DRBD18 appears to play an important role in maintaining life cycle stage identify in *T. brucei*.

## MATERIALS AND METHODS

### Generation of cell lines

PF *T. brucei* strain 29–13 and all cell lines derived from this strain were grown at 27^℃^in SM medium supplemented with 10% fetal bovine serum containing hygromycin (50 µg/mL) and G418 (15 µg/mL). The 29-13-derived procyclic form cell line containing a doxycycline-inducible RNA interference construct that targets the 3’ UTR of DRBD18 (Tb927.11.14090) was described elsewhere (Lott et al. 2015). Endogenous eIF3a subunit was tagged C-terminally with an HA tag by the PCR only methodology (Dean et al. 2015) using forward primer 5’TGCCTACGAAAGGTAAAGTATCTAAGCGTGATGAACAACAAATGCTTCTGGAAATGGAGAAAGAGCGCCTACAAGGGAAGGGTTCTGGTAGTGGTTCC3’ and reverse primer 5’CTACTTATACCACCTATTCCCCTTCAACAGGTACTCACTTATTACCTCTTAAGCTACCGTATTAAGACCCCCTTCGAAGCCCAATTTGAGAGACCTGTGC3’ creating an amplicon with a blasticidin selection cassette. After transfection of parental cells with 10 µg of a gel-purified PCR product, positive transfectants were selected with 20 µg/mL blasticidin and resistant clones were obtained by limiting dilution. To create the KRBP16 3’UTR RNAi cell line, we PCR-amplified a 535-bp fragment of the KRBP16 3’UTR, as annotated in the TriTrypDB gene database (Tb927.11.7900), using *T. brucei* Lister 427 genomic DNA as a template. Forward primer (GATC**GGATCC**CAGTGGTTAAGCGGAGGGGGAAAAAGTTCTTATTCGC) and reverse primer (GATC**AAGCTT**GGAGACACGTTATATATAGCATTAAGACACGCTCAAAAA AAGACGACCTGCACCC) were designed with 5’ overhangs containing the BamHI and HindIII restriction sites, respectively. The KRBP16_3UTR_ RNAi amplicon was blunt-end ligated into the pJet cloning vector using the CloneJET PCR Cloning Kit (ThermoFisher), and the pJet-KRBP16_3UTR_ plasmid was transformed into competent DH5α *E. coli*. The resulting pJet-KRBP16_3UTR_ plasmid was dual-digested with BamHI and HindIII, the KRBP16_3UTR_ insert was gel-purified, and ligated into the p2T7-177 vector in the BamHI and HindIII cloning sites. Approximately 2×10^7^ procyclic 29-13 cells were resuspended in 100 μL of electroporation buffer containing 10 μg of NotI-digested p2T7-177 KRBP16_3UTR_ RNAi plasmid, electroporated using the Lonza Nucleofector device, and immediately transferred to 40 mL of pre-warmed SDM79 growth medium supplemented with 15% FBS. After 24 hours, 1 mL of transfectant parasites were diluted with 19 mL fresh medium, and phleomycin was added to a final concentration of 2.5 μg/mL. Clones were obtained by limiting dilution, and KRBP16 depletion was validated by Western blot.

### Antibodies and western blot

After SDS-PAGE in 12.5% gels, transfer to nitrocellulose membranes, and blocking in TBST with 5% non-fat dry milk, proteins were identified by probing with rabbit polyclonal antibodies against DRBD18 (Lott et al. 2015) (1:2500 dilution), p22 (Hayman et al. 2001) (1:5000 dilution), RBP16 (Hayman et al. 2001) (1:2000 dilution), rat polyclonal antibodies against RBP10 (Wurst et al. 2012) (1:500 dilution; a generous gift from Christine Clayton and Susanne Kramer), or mouse monoclonal antibody against hemagglutinin (Thermo Scientific cat # 26183) (1:5000 dilution). Blots were washed and incubated with either goat anti rabbit HRP or goat anti mouse HRP (1:10000 dilutions). Signal was detected using enhanced chemiluminescence detection (Thermo Scientific, SuperSignal West Pico Plus), imaged in a Chemi Doc system (BioRad), and analyzed using BioRad Image Lab software.

### Polysomal profiles

PF cells were grown to log phase in SMD79 medium at 28°C, and between 1 to 5 × 10^8^ cells were sedimented at 3000 rpm for 10 minutes. Cells were resuspended in 5 mL of medium containing cycloheximide at a final concentration of 100 µg/mL, incubated for 5 minutes, washed in PBS containing cycloheximide, and finally resuspended in 750 µL of buffer A (10 mM Tris pH 7.4, 300 mM KCl, 10 mM MgCl_2_) supplemented with cycloheximide, protease inhibitors (Roche), RNase out (Invitrogen), and 1 mM DTT. Following a 3 minute incubation, 125 µL of lysis buffer (buffer A supplemented with sucrose at 0.2 M final concentration and NP-40 1.2%) were added and the cells were lysed in a Dounce homogenizer (30 to 50 strokes). The lysate was cleared by centrifugation (12,000xg, 10 minutes at 4 C) and supplemented with heparin (10 mg/mL) and RNase out (40U). Equivalent OD260 units of the resulting cytoplasmic extracts were then layered onto 10-40% sucrose linear gradients (generated using a Biocomp gradient maker) and centrifuged at 4 C for 2 hours at 230,000xg using a SW41Ti rotor and an Optima XPN 100 ultracentrifuge. Following ultracentrifugation, the gradient was fractionated using a tube piercer (Brandel) and analyzed on an ISCO UV detector paired to a fraction collector Data were digitized using a DATAQ DI-149 data acquisition instrument. Protein was purified from fractions using methanol chloroform precipitation. Briefly, samples were mixed sequentially with four volumes of methanol, one volume of chloroform, and three volumes of water, and then centrifuged at 14,000xg for one minute. The upper aqueous layer was removed preserving the protein interface which was then washed with four volumes of methanol, and recovered by centrifugation and subsequent drying and resuspension in SDS-PAGE sample buffer.

### Ribosomal profiling

To prepare ribosomal footprints, 5 × 10^8^ PF *T. brucei* cells containing the doxycycline-inducible RNAi construct for DRBD18 silencing were grown in SDM79 medium supplemented with 10% fetal bovine serum at a density of 5-10×10^6^ cells/mL and induced with 4 µg/mL doxycycline for 24 hours. Induced and uninduced cells were then collected by centrifugation at 3000xg for 10 minutes, resuspended in a twentieth of the original volume of medium and subsequently treated with 100 µg/mL cycloheximide for three minutes. The cells were then rapidly chilled rapidly with the addition of 4 volumes ice-cold PBS and centrifuged at 3000xg, 4°C, for 10 minutes, washed in ice-cold polysome buffer (10 mM Tris-HCl pH 7.4, 300 mM KCl and 10mM MgCl_2_) and if necessary quickly frozen in liquid nitrogen. Thawed cells were then resuspended in a twentieth of the volume of ice-cold polysome buffer, and lysed by the addition of ⅓ volume of polysome buffer containing 0.2M sucrose and 0.1%NP-40, incubated 10 minutes on ice and subjected to 30 strokes of a Dounce homogenizer with a tight-inner tolerance pestle. The lysates were cleared by centrifugation at 12.5k rpm for 15 minutes in a microcentrifuge and then treated with RNAse I (1500 U) on ice for 2 hours. Cleared lysates were run on a 10-40% sucrose linear gradient at 4C for 3 hours at 230,000xg. RNA was purified from the monosome fraction (Trizol), run on UREA-PAGE, and size-selected by gel-purifying the 20-100 nt fragment of each lane. The RNA was dephosphorylated with calf intestinal phosphatase, re-precipitated, and was then ready for library preparation. Control, non-ribosome protected RNA was isolated from the cleared lysate (Trizol) and fractionated to a comparable size range by zinc-mediated fragmentation (Invitrogen) prior to gel purification.

### Library preparation, sequencing, and analysis

Libraries were prepared using the Illumina NEBNext Multiplex Small RNA Library Prep Set 1. Samples were sequenced at the UB Genomics and Bioinformatics Core on a HiSeq 2500. Quality filtered reads were mapped to the *T. brucei* TREU927 genome (version 52, http://tritrypdb.org) using bowtie2 (doi.org/10.1038/nmeth.1923) in --very sensitive-local mode. The number of reads per annotated mRNA was determined using featureCounts (doi.org/10.1093/bioinformatics/btt656). The count table was manually curated, and outliers were removed from the analysis. Features with less than 2 counts per million were removed from further analysis (8966 genes passed these criteria). Between 150 and 950 thousand reads per sample were mapped, low counts were due to heavy contamination of rRNA in the sequenced samples. To evaluate differential translation efficiency (TE), the anota2seq package (doi.org/10.1093/nar/gkz223,(Oertlin et al. 2022) was applied with default parameters (log2FC > |1| and FDR less than 0.15). As before, experimental batch effects were introduced as a variable in the linear model. This software classifies mRNAs into 3 categories, the “translation” group is defined as genes where a significant difference in translation occurs because of a change in TE. In the “mRNA abundance” group are mRNAs that significantly change both their translation and mRNA steady-state level (not changing their TE). Finally, the “buffered” group consists of mRNAs that change their steady-state level, but this change does not yield a modification of the translation level of the protein (by modification of the TE). Overrepresentation of GO terms among DEGs lists was established using the tools available at TritrypDB (http://tritrypdb.org/) using an adjusted p-value of less than 0.05 as a cutoff for significance. Statistical analysis and plots were performed in R, unless otherwise specified. Python in-house scripts were used to analyze the size of mapped reads in each sample.

### Proximity Ligation Assay

PF were fixed with 2% formaldehyde, deposited on poly-L-lysine coated slides, permeabilized with 0.5% NP40, and blocked with Duolink® Blocking Solution (SIGMA) for 60 minutes at 37C. The fixed cells were first incubated with primary antibodies against target proteins raised in different species (rabbit anti-DRBD18 (Lott et al. 2015) and mouse monoclonal anti-HA, cat # 26183, Invitrogen) and then with Duolink® Probes MINUS and PLUS. The probes were ligation and amplified according to manufacturer’s instructions. Serial image stacks (0.2 micron Z-increment) were collected with capture times from 100–400 msec (100x PlanApo, oil immersion, 1.46 na) on a motorized Zeiss Axioimager M2 stand equipped with a rear-mounted excitation filter wheel, a triple pass (DAPI/FITC/Texas Red) emission cube, differential interference contrast (DIC) optics, and an Orca ER CCD camera (Hamamatsu, Bridgewater, NJ). Images were collected using Volocity 6.1 Acquisition Module (Improvision Inc., Lexington, MA) and individual channel stacks were deconvolved by a constrained iterative algorithm, pseudocolored, and merged using Volocity 6.1 Restoration Module.

### Proteomics

*Sample preparation*. A surfactant cocktail-aided extraction/precipitation/on-pellet digestion (SEPOD) protocol was used for sample preparation as previously described. (Shen et al. 2018a) *T. brucei* cell pellets were suspended in 400 μL ice-cold surfactant cocktail buffer (50 mM Tris-formic acid, 150 mM NaCl, 2% SDS, 0.5% sodium deoxycholate, 2% IGEPAL CA630, pH 8.4) supplemented with cOmplete protease inhibitor cocktail tablets (Roche Applied Science, Indianapolis, IN, USA). Samples were vortexed, placed on ice for 30 min for cell lysis, and were sonicated by 3 sonication-cooling cycles (10 s each) using a high-energy probe sonicator (Qsonica, Newtown, CT, USA). Protein lysates were centrifuged at 18,000 x g, 4 °C for 30 min, and supernatant were transferred to new Eppendorf tubes. Protein concentration was determined by bicinchoninic acid assay. For protein digestion, 100 μg protein was aliquoted from each sample. Protein was reduced by 10 mM dithiothreitol (DTT) at 56 °C for 30 min and then alkylated by 25 mM iodoacetamide (IAM) at 37 °C for 30 min in darkness. Both steps were performed with constant shaking at 550 rpm in a covered thermomixer (Eppendorf, Framingham, MA, USA). Protein was precipitated by adding 6 volumes of ice-cold acetone with vigorous vortexing, and the mixture was incubated at -20 °C for 3 h. Protein precipitated was pelleted by centrifugation at 18,000 x g, 4 °C for 30 min, and was gently rinsed by 500 μL methanol. After removing all liquid, protein pellet was left to air dry for 1 min, and 80 μL Tris-formic acid (FA) pH 8.4 was added to wet the pellet. A total volume of 20 μL trypsin (Sigma-Aldrich, St. Louis, MO, USA) dissolved in 50 mM Tris-FA pH 8.4 (0.25 μg/ μL) was added to each sample to reach a final enzyme-to-substrate ratio of 1:20 (w/w), and samples were incubated in a covered thermomixer at 37 °C for 6h with constant shaking. Tryptic digestion was terminated by addition of 1 μL FA, and samples were centrifuged at 18,000 x g, 4 °C for 30 min. Supernatant was carefully transferred to LC vials for analysis.

*LC-MS analysis*. The LC-MS system consists of a Dionex UltiMate 3000 nano LC system, a Dionex UltiMate 3000 micro LC system with a WPS-3000 autosampler, and an Orbitrap Fusion Lumos mass spectrometer (ThermoFisher Scientific, San Jose, CA, USA). A large-inner diameter (i.d.) trapping column (300-μm i.d. x 5 mm, Agilent Technologies, Santa Clara, CA, USA) was coupled to the analytical column (75-μm i.d. x 65 cm, packed with 2.5-μm XSelect CSH C18 material) for high-capacity sample loading, cleanup and delivery. For each sample, peptide equivalent to 4 μg protein was injected for LC-MS analysis. Mobile phase A and B were 0.1% FA in 2% acetonitrile (ACN) and 0.1% FA in 88% ACN. The 180-min LC gradient profile for the analytical column was: 4% B for 3 min, 4-9% B for 5 min, 9-30% B for 117 min, 30-50% B for 10 min, 50-97% B for 1 min, 97% B for 17 min, and then equilibrated to 4% for 27 min. Mass spectrometer was operated under data-dependent acquisition (DDA) mode with a maximal duty cycle of 3 s. MS1 spectra was acquired by Orbitrap (OT) under 240k resolution for ions in the m/z range of 400-1,500. Automatic Gain Control (AGC) target and maximal injection time was set to 175% and 50 ms, and dynamic exclusion was set as 60 s, ± 10 ppm. Precursor ions were isolated by quadrupole with a 1.6-Th m/z window for fragmentation by high-energy collisional dissociation (HCD) at 30% energy. MS2 spectra were acquired by OT under 15k resolution. AGC target and maximal injection time was set to 100% and 22 ms. Detailed LC-MS settings and information can be found in our previous publications (Shen et al. 2017; Shen et al. 2018b; Wang et al. 2021).

*Data processing and analysis*. An in-house developed UHR-IonStar pipeline was employed for data processing and analysis. For protein identification, database searching was performed using the MS-GF+ search engine (v20210108, released in January 2021). Search parameters included: 1) Protein database: human Swiss-Prot protein sequence database (20,302 entries, downloaded in May 2020); 2) Precursor mass tolerance: 20 ppm; 3) Instrument type: Q-Exactive; 4) Matches per spectrum:1; 5) Dynamic modifications: oxidation of Methionines (M) and acetylation of peptide N-termini; 6) Fixed modification: carbamidomethylation of Cysteines (C); 7) Maximal missed cleavages: 2. Peptide-spectrum match (PSM) filtering, protein inference/grouping, and false discovery rate (FDR) control were performed by IDPicker (v3.1.18192.0). Protein/peptide FDR was set to 1%, and a minimum of 2 unique peptides per protein was set. Proteins with no unique peptides were grouped with a maximum of 50 proteins per protein group. The filtered PSM list was generated by the UHR-IonStar APP (https://github.com/JunQu-Lab/UHRIonStarApp) using protein/peptide/PSM lists exported from IDPicker. For protein quantification, peptide quantitative features were extracted from LC-MS files using SIEVE (v2.2, Thermo Fisher Scientific, San Jose, CA, USA) and processed by the UHR-IonStar APP to generate final quantification results. Main procedures included: 1) Chromatographic alignment with ChromAlign (Sadygov et al. 2006) for dataset-wide retention time (RT) calibration and peak clustering. Performance of the alignment step was evaluated by alignment scores in the entire dataset; 2) Data-independent MS1 quantitative feature extraction using the Direct Ion-Current Extraction (DICE) method, which utilizes a pre-defined m/z-RT window (10 ppm, 1 min for 240 MS1 acquisition) to extract ion chromatograms for all precursor ions subjected to fragmentation and MS2 acquisition in the aligned dataset. Each set of ion chromatograms with corresponding area under the curve (AUC) in the dataset was termed as a “frame”; 3) Integration of the SIEVE frame database and the filtered PSM list by a unique identifier combining file name and MS2 scan number. Frames with valid peptide sequences were subjected to frame-level quality control, global data normalization, peptide-level outlier detection, and aggregation to protein level. More detailed information about the UHR-IonStar pipeline can be found in our previous publications (Shen et al. 2018b; Wang et al. 2021). Quantification results were further processed by the UHR-IonStar APP for data formatting/cleanup, statistical testing by Student’s t-test, and inter-group protein ratio calculation.

### Data availability

RNA sequencing results are available at the Sequence Read Archive under project number PRJNA913808. The mass spectrometry proteomics data have been deposited to the ProteomeXchange Consortium via the PRIDE (Perez-Riverol et al. 2022) partner repository with the dataset identifier PXD039064.

## Acknowledgements

We thank Christine Clayton and Susanne Kramer for anti-RBP10 antibodies, Parul Pandey for critical reading of the manuscript, and Brianna Tylec for assistance with figure preparation. We also thank the UB Genomics and Bioinformatics Core, especially Don Yergeau. This work was supported by NIH R01AI141557 to LKR. PS and JSS received financial support from PEDECIBA and are members of ANII research career.

## SUPPLEMENTAL FIGURE AND TABLE LEGENDS

**Figure S1. RBP16-depleted cells exhibit a growth defect but no change in monosome/polysome ratio**. (*A*) Growth of procyclic DRBD18 RNAi cells and RPB16 RNAi cells, either uninduced or induced with doxycycline. (*B*) Western blots showing degree of knockdown. *(C)* Control showing that depletion of an essential protein unrelated to DRBD18 (RBP16) does not result in an early change the polysome profile, demonstrating that the increased monosome/polysome ratio observed in DRBD18 knockdown cells is not due to indirect cell death effects.

**Figure S2. Size distribution of the reads from ribosomal footprint samples and total RNA controls**. Histograms showing length distributions of reads and their abundances in samples obtained using the ribosomal footprint protocol (left panels, samples S01, S02, S05, S06, S09, S10) compared to the total RNA protocol (right panels; samples S03, S04, S07, S08, S11, S12). All cell counts were normalized so that the area under the curve equals 1.

**Figure S3. Transcripts that are translationally repressed after DRBD18 depletion normally have higher abundance than those that are translationally enhanced**. Available datasets (Kolev et al. 2010) were used to plot the values of transcript abundance (mRNA copies per cell) for each gene in the TE UP or TE DOWN categories.

**Table S1. List of ribosomal proteins and eIF3 subunits identified by MS bound to DRBD18-TAP** (Lott et al. 2015).

**Table S2. Transcripts whose TE is affected by DRBD18 knockdown**.

**Table S3. Anota2seq analysis of transcripts with altered translation efficiency, buffered translation, or altered abundance in induced vs. uninduced DRBD18 knockdown cells**.

**Table S4. Changes in the proteome in doxycycline-induced PF DRBD18 knockdown cells**. Dysregulated proteins are defined as those with p-value <0.05 and fold change >50% (*i.e*., protein ratio >1.5 or <0.67).

## REFERENCES

Archer SK, Luu VD, de Queiroz RA, Brems S, Clayton C. 2009. Trypanosoma brucei PUF9 regulates mRNAs for proteins involved in replicative processes over the cell cycle. PLoS Pathog 5: e1000565.

Bishola Tshitenge T, Clayton C. 2022. The Trypanosoma brucei RNA-binding protein DRBD18 ensures correct mRNA trans splicing and polyadenylation patterns. RNA 28: 1239–1262.

Chassé H, Boulben S, Costache V, Cormier P, Morales J. 2017. Analysis of translation using polysome profiling. Nucleic Acids Res 45: e15.

Chavez S, Urbaniak MD, Benz C, Smircich P, Garat B, Sotelo-Silveira JR, Duhagon MA. 2021. Extensive translational regulation through the proliferative transition of Trypanosoma cruzi revealed by multi-omics. mSphere 6: e0036621.

Christiano R, Kolev NG, Shi H, Ullu E, Walther TC, Tschudi C. 2017. The proteome and transcriptome of the infectious metacyclic form of Trypanosoma brucei define quiescent cells primed for mammalian invasion. Mol Microbiol 106: 74–92.

Clayton C. 2019. Regulation of gene expression in trypanosomatids: living with polycistronic transcription. Open Biol 9: 190072.

De Pablos LM, Kelly S, de Freitas Nascimento J, Sunter J, Carrington M. 2017. Characterization of RBP9 and RBP10, two developmentally regulated RNA-binding proteins in Trypanosoma brucei. Open Biol 7:160159.

Dean S, Sunter J, Wheeler RJ, Hodkinson I, Gluenz E, Gull K. 2015. A toolkit enabling efficient, scalable and reproducible gene tagging in trypanosomatids. Open Biol 5: 140197.

Dean S, Sunter JD, Wheeler RJ. 2017. TrypTag.org: A trypanosome genome-wide protein localisation resource. Trends Parasitol 33: 80–82.

Durante IM, Butenko A, Rašková V, Charyyeva A, Svobodová M, Yurchenko V, Hashimi H, Lukeš J. 2020. Large-scale phylogenetic analysis of trypanosomatid adenylate cyclases reveals associations with extracellular lifestyle and host–pathogen interplay. Genome Biology and Evolution 12: 2403–2416.

El Mouali Y, Balsalobre C. 2019. 3’ untranslated regions: regulation at the end of the road. Curr Genet 65: 127–131.

Fredriksson S, Gullberg M, Jarvius J, Olsson C, Pietras K, Gustafsdottir SM, Ostman A, Landegren U. 2002. Protein detection using proximity-dependent DNA ligation assays. Nat Biotechnol 20: 473–477.

Grant CE, Bailey TL. 2021. XSTREME: comprehensive motif analysis of biological sequence datasets. BioRxiv: https://doi.org/10.1101/2021.1109.1102.458722.

Gunasekera K, Wuthrich D, Braga-Lagache S, Heller M, Ochsenreiter T. 2012. Proteome remodelling during development from blood to insect-form Trypanosoma brucei quantified by SILAC and mass spectrometry. BMC Genomics 13: 556.

Hayman ML, Miller MM, Chandler DM, Goulah CC, Read LK. 2001. The trypanosome homolog of human p32 interacts with RBP16 and stimulates its gRNA binding activity. Nucleic Acids Res 29: 5216–5225.

Ingolia NT, Ghaemmaghami S, Newman JR, Weissman JS. 2009. Genome-wide analysis in vivo of translation with nucleotide resolution using ribosome profiling. Science 324: 218–223.

Jensen BC, Ramasamy G, Vasconcelos EJ, Ingolia NT, Myler PJ, Parsons M. 2014. Extensive stage-regulation of translation revealed by ribosome profiling of Trypanosoma brucei. BMC Genomics 15: 911.

Johansson J, Freitag NE. 2019. Regulation of Listeria monocytogenes virulence. Microbiol Spectr 7(4).

Klein C, Terrao M, Clayton C. 2017. The role of the zinc finger protein ZC3H32 in bloodstream-form Trypanosoma brucei. PLoS One 12: e0177901.

Klein C, Terrao M, Inchaustegui Gil D, Clayton C. 2015. Polysomes of Trypanosoma brucei: Association with initiation factors and RNA-binding proteins. PLoS One 10: e0135973.

Kolev NG, Franklin JB, Carmi S, Shi H, Michaeli S, Tschudi C. 2010. The transcriptome of the human pathogen Trypanosoma brucei at single-nucleotide resolution. PLoS Pathog 6: e1001090.

Kolev NG, Ullu E, Tschudi C. 2014. The emerging role of RNA-binding proteins in the life cycle of Trypanosoma brucei. Cellular Microbiology 16: 482–489.

Kramer S, Carrington M. 2011. Trans-acting proteins regulating mRNA maturation, stability and translation in trypanosomatids. Trends Parasitol 27: 23–30.

Li K, Zhou S, Guo Q, Chen X, Lai DH, Lun ZR, Guo X. 2017. The eIF3 complex of Trypanosoma brucei: composition conservation does not imply the conservation of structural assembly and subunits function. RNA 23: 333–345.

Lopez MA, Saada EA, Hill KL. 2015. Insect stage-specific adenylate cyclases regulate social motility in African trypanosomes. Eukaryot Cell 14: 104–112.

Lott K, Mukhopadhyay S, Li J, Wang J, Yao J, Sun Y, Qu J, Read LK. 2015. Arginine methylation of DRBD18 differentially impacts its opposing effects on the trypanosome transcriptome. Nucleic Acids Res 43: 5501–5523.

Matthews KR. 2005. The developmental cell biology of Trypanosoma brucei. Journal of Cell Science 118: 283–290.

McDonald L, Cayla M, Ivens A, Mony BM, MacGregor P, Silvester E, McWilliam K, Matthews KR. 2018. Non-linear hierarchy of the quorum sensing signalling pathway in bloodstream form African trypanosomes. PLoS Pathog 14: e1007145.

Mishra A, Kaur JN, McSkimming DI, Hegedusova E, Dubey AP, Ciganda M, Paris Z, Read LK. 2021. Selective nuclear export of mRNAs is promoted by DRBD18 in Trypanosoma brucei. Mol Microbiol 116: 827–840.

Mugo E, Clayton C. 2017. Expression of the RNA-binding protein RBP10 promotes the bloodstream-form differentiation state in Trypanosoma brucei. PLoS Pathog 13: e1006560.

Oertlin C, Watt K, Ristau J, Larsson O. 2022. Anota2seq analysis for transcriptome-wide studies of mRNA Ttanslation. Methods Mol Biol 2418: 243–268.

Ojo KK, Gillespie JR, Riechers AJ, Napuli AJ, Verlinde CLMJ, Buckner FS, Gelb MH, Domostoj MM, Wells SJ, Scheer A et al. 2008. Glycogen synthase kinase 3 is a potential drug target for African trypanosomiasis therapy. Antimicrob Agents Chemother 52: 3710–3717.

Pelletier M, Read LK. 2003. RBP16 is a multifunctional gene regulatory protein involved in editing and stabilization of specific mitochondrial mRNAs in Trypanosoma brucei. RNA 9: 457–468.

Perez-Riverol Y, Bai J, Bandla C, Garcia-Seisdedos D, Hewapathirana S, Kamatchinathan S, Kundu DJ, Prakash A, Frericks-Zipper A, Eisenacher M et al. 2022. The PRIDE database resources in 2022: a hub for mass spectrometry-based proteomics evidences. Nucleic Acids Res 50: D543–D552.

Quintana JF, Zoltner M, Field MC. 2021. Evolving differentiation in African trypanosomes. Trends Parasitol 37: 296–303.

Rico-Jimenez M, Ceballos-Perez G, Gomez-Linan C, Estevez AM. 2021. An RNA-binding protein complex regulates the purine-dependent expression of a nucleobase transporter in trypanosomes. Nucleic Acids Res 49: 3814–3825.

Rowe W, Kershaw CJ, Castelli LM, Costello JL, Ashe MP, Grant CM, Sims PFG, Pavitt GD, Hubbard SJ. 2014. Puf3p induces translational repression of genes linked to oxidative stress. Nucleic Acids Res 42: 1026–1041.

Sadygov RG, Maroto FM, Huhmer AF. 2006. ChromAlign: A two-step algorithmic procedure for time alignment of three-dimensional LC-MS chromatographic surfaces. Anal Chem 78: 8207–8217.

Salmon D. 2018. Adenylate cyclases of Trypanosoma brucei, environmental sensors and controllers of host innate immune response. Pathogens 7: 48.

Shaw S, Knusel S, Abbuhl D, Naguleswaran A, Etzensperger R, Benninger M, Roditi I. 2022. Cyclic AMP signalling and glucose metabolism mediate pH taxis by African trypanosomes. Nat Commun 13: 603.

Shen S, An B, Wang X, Hilchey SP, Li J, Cao J, Tian Y, Hu C, Jin L, Ng A et al. 2018a. Surfactant cocktail-aided extraction/precipitation/on-pellet digestion strategy enables efficient and reproducible sample preparation for large-scale quantitative proteomics. Anal Chem 90: 10350–10359.

Shen X, Shen S, Li J, Hu Q, Nie L, Tu C, Wang X, Orsburn B, Wang J, Qu J. 2017. An IonStar experimental strategy for MS1 ion current-based quantification using ultrahigh-field Orbitrap: Reproducible, in-depth, and accurate protein measurement in large cohorts. J Proteome Res 16: 2445–2456.

Shen X, Shen S, Li J, Hu Q, Nie L, Tu C, Wang X, Poulsen DJ, Orsburn BC, Wang J et al. 2018b. IonStar enables high-precision, low-missing-data proteomics quantification in large biological cohorts. Proc Natl Acad Sci U S A 115: E4767–E4776.

Smircich P, Eastman G, Bispo S, Duhagon MA, Guerra-Slompo EP, Garat B, Goldenberg S, Munroe DJ, Dallagiovanna B, Holetz F et al. 2015. Ribosome profiling reveals translation control as a key mechanism generating differential gene expression in Trypanosoma cruzi. BMC Genomics 16: 443.

Tinti M, Ferguson MAJ. 2022. Visualisation of proteome-wide ordered protein abundances in. Wellcome Open Res 7: 34.

Valasek LS, Zeman J, Wagner S, Beznoskova P, Pavlikova Z, Mohammad MP, Hronova V, Herrmannova A, Hashem Y, Gunisova S. 2017. Embraced by eIF3: structural and functional insights into the roles of eIF3 across the translation cycle. Nucleic Acids Res 45: 10948–10968.

Vasquez JJ, Hon CC, Vanselow JT, Schlosser A, Siegel TN. 2014. Comparative ribosome profiling reveals extensive translational complexity in different Trypanosoma brucei life cycle stages. Nucleic Acids Res 42: 3623–3637.

Verma-Gaur J, Traven A. 2016. Post-transcriptional gene regulation in the biology and virulence of Candida albicans. Cell Microbiol 18: 800–806.

Wang X, Jin L, Hu C, Shen S, Qian S, Ma M, Zhu X, Li F, Wang J, Tian Y et al. 2021. Ultra-high-resolution IonStar strategy enhancing accuracy and precision of MS1-based proteomics and an extensive comparison with state-of-the-art SWATH-MS in large-cohort quantification. Anal Chem 93: 4884–4893.

Wolf DA, Lin Y, Duan H, Cheng Y. 2020. eIF-Three to Tango: emerging functions of translation initiation factor eIF3 in protein synthesis and disease. J Mol Cell Biol 12: 403–409.

Wurst M, Seliger B, Jha BA, Klein C, Queiroz R, Clayton C. 2012. Expression of the RNA recognition motif protein RBP10 promotes a bloodstream-form transcript pattern in Trypanosoma brucei. Mol Microbiol 83: 1048–1063.

